# Modulatory effect of plasma-activated water on arbuscular mycorrhizal symbiosis in *Lotus japonicus*

**DOI:** 10.1101/2024.11.19.624245

**Authors:** Enrico Cortese, Filippo Binci, Erfan Nouri, Arianna Capparotto, Alessio G. Settimi, Manuele Dabalà, Vanni Antoni, Andrea Squartini, Marco Giovannetti, Lorella Navazio

**Affiliations:** Department of Biology, University of Padova, Via U. Bassi 58/B, 35131 Padova, Italy; Department of Agronomy, Food, Natural Resources, Animals and Environment (DAFNAE), University of Padova, Viale dell’Università 16, 35020 Legnaro, Italy; Department of Industrial Engineering, University of Padova, Via Marzolo 9, 35131 Padova, Italy; National Research Council, Institute for Plasma Science and Technology (CNR-ISTP), Corso Stati Uniti 4, 35127 Padova, Italy; Consorzio RFX, Corso Stati Uniti 4, 35127 Padova, Italy

**Keywords:** Arbuscular mycorrhizal symbiosis, Calcium signalling, Plasma-activated water, Plasma agriculture, *Lotus japonicus*

## Abstract

Plasma-activated water (PAW) is a recently developed cutting-edge technology that is increasingly gaining interest for its applications in medicine, food industry and agriculture. In plant biology, PAW has been shown to promote seed germination, plant growth, and plant resistance to biotic and abiotic stresses. Despite increasing knowledge of the beneficial effects exerted by PAW on plants, little information is currently available about how this emerging technology may affect the mutualistic plant-microbe interactions in the rhizosphere. In this work we have investigated the impact of irrigation with PAW, generated by a plasma torch, on arbuscular mycorrhizal (AM) symbiosis between the model legume *Lotus japonicus* and the AM fungus *Rhizophagus irregularis.* Since PAW sensing by plants has recently been demonstrated to occur through calcium-mediated signalling, we monitored early cellular responses to different doses of PAW in *L. japonicus* roots expressing the Ca^2+^-sensitive photoprotein aequorin targeted to either the cytosol or nucleus. Quantitative analyses of AM fungal accommodation in host roots along with phosphate accumulation in leaves, as well as chemical analysis of N, C, S in shoots, showed that treatments with PAW play a modulatory role on plant AM symbiotic performance, in a manner dependent on the time interval of water exposure to the plasma and on the duration of plant irrigation treatment with PAW. Establishing a solid scientific ground for plasma-related technology may provide key elements to develop tools and treatments aimed to increase crop plant yield in a sustainable manner.

## 1. Introduction

The increasing global demand for food, combined with the limits on natural resources and the issues related to the impact of food production on the environment, are fostering the quest for new technologies in agriculture and food systems in general. Plasma-activated water (PAW) is a recently emerged promising technology, which may contribute to alleviate the aforementioned issues. PAW is generated by exposing water (or other liquids) to atmospheric non-thermal plasma, i.e. an ionized gas characterized by relatively weak ionization with electrons at higher temperature than heavy particles (e.g. ions, molecules, radicals). The condition of non-equilibrium favours the presence of excited chemicals which, when interacting with water, result in a wide range of reactive oxygen species (ROS) and reactive nitrogen species (RNS) with varying half-lives (Fridman & Kennedy, 2021). The rich mixture of chemicals has been shown to stimulate biological responses when applied to living systems (Kaushik et al., 2018; Thirumdas et al., 2018). In particular, the resulting PAW is increasingly seen as a safe, rather inexpensive, and eco-friendly alternative that may reduce the use of pesticides and fertilizers in agriculture, thanks to its antimicrobial/disinfection properties as well as to its effects on the improvement of seed germination and plant growth. Moreover, the ability of PAW to mildly induce plant defence responses, effectively boosting plant resistance against subsequent pathogen attacks (a pre-alert state termed “priming”) is leading the research towards the fine-tuning of this novel “green” technology to maximize its beneficial effects in the context of a more sustainable agriculture, according to the strategy “from Farm to Fork” (Adhikari et al., 2020; Holubová et al., 2020; Pańka et al., 2022).

A relevant subject of potential great impact, but rather unexplored so far, is how PAW treatment may impact beneficial plant-microbe interactions in the rhizosphere. The arbuscular mycorrhizal (AM) symbiosis, established between most land plants and Glomeromycotina fungi (Genre et al., 2020), and the symbiotic nitrogen fixation, a mutualistic interaction between legumes and rhizobia (Gage, 2004), positively influence plant growth, by improving plant mineral nutrition and increasing plant tolerance to abiotic and biotic stresses. In addition, they offer ecosystem services in natural and agricultural environments. In the current context of growing environmental concerns, AM and nitrogen-fixing symbioses play a key role in ecosystem functioning and environment restoration. The double symbiotic aptitude in staple leguminous crops raises great interest in sustainable agriculture to feed the growing global population. Moreover, increasing threats to crops by the ongoing climate change and their consequences on soil nutrient and water availability made the development of biostimulants of plant-microbe symbioses an urgent strategy to strengthen plant tolerance to abiotic stresses and enhance plant development (Trivedi et al., 2022). As these symbiotic associations are crucial to overall plant health and productivity, understanding how PAW and in general plasma-based treatments may affect them is fundamental for large scale agricultural implementation of this innovative technology. It has recently been demonstrated that pre-sowing treatment of soybean and pea seeds with non-thermal plasmas, generated by a dielectric barrier discharge source, resulted in the enhancement of nodulation and biological nitrogen fixation (Pérez-Pizá et al., 2020, Abeysingha et al., 2024). Moreover, novel approaches for industrial-scale nitrogen fixation based on atmospheric plasma, implying lower energy consumption than the traditional Haber-Bosch process, are currently under investigation (Aceto et al., 2024).

In this work we investigated how *Lotus japonicus* plants respond to treatment with PAW generated by a plasma torch, in terms of early Ca^2+^-mediated root responses and establishment and development of the AM symbiosis. The obtained data provide insights into the potential use of PAW to improve plant mineral nutrition and further develop sustainable agricultural practices.

## 2. Materials and Methods

### 2.1 Sterilization and germination of Lotus japonicus seeds

*Lotus japonicus* Gifu ecotype seeds were scarified in a mortar with sandpaper. The sterilization was performed in a 2-mL tube with 2 mL of 0.5% NaClO for 11 min. Seeds were washed five times with sterile H_2_O and left swollen in water for about 30 minutes. Seeds were plated in half-strength B5 (Duchefa Biochemie, Haarlem, The Netherlands) medium (pH 5.5, adjusted with KOH), supplemented with 1% (w/v) Plant Agar (Duchefa) in 12×12 cm squared Petri dishes and were vertically incubated covered with an aluminum foil. After 3 days, the aluminum foil was removed and seedlings were left growing in the same conditions.

### 2.2 L. japonicus hairy root transformation by Agrobacterium rhizogenes

Hairy roots transformation was performed according to Boisson-Dernier et al. (2001). −80°C stocked *Agrobacterium rhizogenes* AR1193 strains carrying the plasmid encoding either cytosolic or nuclear-targeted aequorin-based Ca^2+^ chimeras (Binci et al., 2024) were plated in LB medium containing 1% Bacto-Agar and supplemented with 50 µg/ml rifampicin, 100 µg/ml ampicillin and 50 µg/ml kanamycin. Bacteria were grown for 2 days at 28°C, and then plated again to have a full Petri dish of actively growing cells. 7-day-old *L. japonicus* seedlings were cut with a scalpel in the hypocotyl region to remove the root. The scar at the base of the remaining shoot was then dipped into the bacterial film on the plate and placed in a new squared Petri plate provided with half-strength B5, 1% Plant Agar. The shoots were then co-cultivated with bacteria at 22°C for 3 days in the dark and for additional 3 days at 16h/8h light/dark cycle. The shoots were then transferred to new squared 12×12 cm Petri dishes containing half-strength B5, 1% Plant Agar and 300 µg/ml cefotaxime (Duchefa) and let grow for 4-5 weeks (22°C, 50% humidity, 16h/8h light/dark). Transformed roots were then identified at the fluorescence stereomicroscope MZ16f (Leica, Wetzlar, Germany) via the fluorescence of the YFP-aequorin chimeras (Binci et al., 2024).

### 2.3 Generation of plasma-activated water (PAW)

Plasma-activated water (PAW) was produced by exposing deionized H_2_O to the atmospheric plasma generated by a plasma torch operating at 900 W power and 3 bar pressure. The torch used for this study was a single rotating FLUME Jet RD1004 with an FG 1001 plasma generator (Plasmatreat, Elgin, IL, USA) with a maximum power of 2.7 kW (230 V, 12 A). Throughout the study, the reported results employed 50 mL of deionized H_2_O with its surface exposed for 5 to 10 min at 1.5 cm from the torch nozzle. H_2_O was kept in beakers immersed in an ice and salt cooling bath (Supplementary Fig. S1) to keep the water temperature constat during the exposure. Once generated, PAW was divided into single-use aliquots, rapidly cryogenically frozen through immersion in liquid nitrogen, and stored at −80°C.

### 2.4 Evaluation of cell viability

Cell viability was assessed using the Evans blue method (Baker & Mock, 1994) using *L. japonicus* suspension cell cultures, established and subcultured as previously described (Moscatiello *et al*., 2018). Cultured-cells were either maintained in control conditions or treated with PAW (diluted 1:2 and 1:4) for 1 h or 48 h during their mid-exponential growth phase (4 days). Following a 15 min incubation with 0.05% (w/v) Evans blue dye (Merck, Darmstadt, Germany), unbound dye was removed by thorough washing with H_2_O. The dye bound to dead cells was then solubilized in a solution containing 1% (w/v) SDS and 50% (v/v) methanol, maintained at 55°C for 30 min and then collected. The extent of cell death was determined by measuring the absorbance at 600 nm. As positive control representing 100% cell death, a separate cell aliquot was incubated at 100°C for 10 min.

### 2.5 Aequorin-based Ca^2+^ measurement assays

Ca^2+^ measurement assays were conducted in *L. japonicus* roots using the recombinant expression of aequorin chimeras targeted to either the cytosol or nucleus (Binci et al., 2024). 5-mm-long transformed root segments from composite plants were reconstituted overnight with 5 µM coelenterazine (Prolume, Pinetop, AZ, USA). On the following day, the root segments were extensively washed to remove excess coelenterazine and transferred in a custom-built luminometer (ET Enterprises Ltd, Uxbridge, UK) containing a 9893/350A photomultiplier (Thorn EMI). Each root segment was placed in 50 µl H_2_O and challenged by injection of an equal volume of PAW produced through a 5 min long exposure to plasma torch at different dilutions (1:2, 1:4, 1:8). Ca^2+^ dynamics were recorded for 20 min and each experiment was terminated with the injection of 100 µL of the discharge solution (30% v/v ethanol, 1 M CaCl_2_). The light signal was recorded and later converted into Ca^2+^ concentration values using an algorithm based on the Ca^2+^ response curve of aequorin (Ottolini et al., 2014).

### 2.6 PAW treatment of L. japonicus seedlings co-cultivated with the AM fungus Rhizophagus irregularis

14 days after sterilization, *L. japonicus* seedlings were transplanted into 9×9 cm plastic pots with 4 seedlings per pot. The pots were filled with washed and autoclaved river sand characterized by a granulometry of 1-5 mm, which was subsequently inoculated with 3000 propagules of the AM fungus *Rhizophagus irregularis* (Agronutrition, Carbonee, France). Afterwards, each pot was irrigated using 20 ml of a modified Long Ashton liquid medium (1.5 mM Ca(NO_3_)_2_, 1 mM KNO_3_, 0.75 mM MgSO_4_, 0.1 mM Fe-EDTA, 10 µM MnCl_2_, 50 µM H_3_BO_3_, 1.75 µM ZnCl_2_, 0.5 µM CuCl_2_, 0.8 µM Na_2_MoO_4_, 1 µM KI, 0.1 µM CoCl_2_) with low phosphate content (20 µM KH_2_PO_4_). Starting from the 10^th^ day of pot cultivation, a bi-weekly irrigation regimen was initiated. The first irrigation involved applying 20 ml of the appropriate treatment, either H_2_O (control samples) or freshly prepared PAW activated by plasma torch for 5 min (PAW 5’) or 10 min (PAW 10’) (Supplementary Fig. S2). The second irrigation, conducted 3 days later in the same week, employed 20 ml of modified liquid Long Ashton, containing 20 µM Pi. The duration of the experiments varied between 4 and 7 weeks in order to assess the impact of PAW treatment at distinct stages of the AM symbiosis establishment and functioning. All irrigations were performed from the bottom of the pot. At the end of each irrigation period *L. japonicus* plants were harvested by pooling together the 4 seedlings growing in each pot, representing a single biological replicate. The roots were separated from the shoots of each plant and rinsed gently with H_2_O to remove excess sand debris. For each plant, the fresh biomasses of the root apparatus, the entire shoot and the second leaf triplet from the shoot apex were evaluated with a precision digital balance. For each plant, roots and aerial parts (either entire shoots or only leaves) were processed as described in the next sections.

### 2.7 Evaluation of AM fungal colonization

Roots of inoculated plants were ink-stained to highlight intraradical fungal structures and quantify them according to the method described by Trouvelot et al. (1986). Before staining, the roots were clarified with 10% KOH at 96°C on a heating block for 6 min. After incubation, KOH was discarded and 2 washing steps were performed with 5% lactic acid. Roots were then incubated in the staining solution (5% black ink, 5% lactic acid) at 96°C for 11 minutes. The staining solution was then discarded and stained roots were washed in 5% lactic acid, replacing the washing solution until the liquid was clear. From each biological replicate (1 pot, 4 plants) five microscope slides were prepared, each containing 20 root fragments of 1 cm in length. Consequently, for each pot approximately 100 cm of root apparatus were analysed. The level of mycorrhizal colonization was assessed according to the Trouvelot method (Trouvelot et al., 1986).

### 2.8 Measurements of phosphate concentration

Each leaf sample was weighted in an analytical balance and stored at −80°C until use. Frozen samples were homogenized with the help of a metal bead in a TissueLyser II (Qiagen, Hilden, Germany) via two rounds of shaking at 30 Hz for 20 s. The tissue powder was resuspended in 1 ml ultrapure H_2_O, vortexed for 10 s and incubated at 100°C for 1 h. After 10 min incubation on ice, samples were centrifuged at 4°C for 15 min at 13200 rpm. The supernatant was used for measurement of the concentration of soluble phosphate using the Phosphate Assay Kit (Merck, Darmstadt, Germany), following the manufacturer’s instructions. Standards were prepared to make a calibration curve ranging from 0 to 40 µM phosphate. Both standards and samples were mixed with an equal volume of the dye Malachite green in a transparent 96-well plate and incubated for 30 min at room temperature in the dark. Absorbance at 620 nm was measured using a Tecan multiwell plate reader (Männedorf, Switzerland). Each sample was analysed in two technical replicates. Absorbance data were converted into concentrations of phosphate using the calibration curve and standardized to the fresh weight of the leaf sample.

### 2.9 Chemical analyses of L. japonicus shoot samples

Shoots from harvested seedlings were used to quantify N, C and S concentrations. To this aim, the samples were dried in an oven set at 60°C until a constant weight was achieved (2-16 h). The dried plant samples were ground into a fine powder to ensure homogeneity. Chemical analyses of N, C and S were conducted using a CNS Macrovario combustion analyzer (Macrovario, GmbH).

## 3. Results

### 3.1 Aequorin-based Ca^2+^ measurements in Lotus japonicus roots in response to PAW

Plasma-activated water (PAW), generated by exposure of water to different plasma sources and conditions to span a wide range of doses and chemical mixtures, has recently been shown to induce rapid and remarkable elevations in the intracellular concentration of Ca^2+^ in the model plant *Arabidopsis thaliana* (Cortese et al., 2021 and 2022). Unravelling the mechanisms of perception of PAW by plants may offer a solid scientific background to modulate the large variety of effects that PAW play on plants, ranging from the promotion of plant growth to increased resistance to environmental stresses (Holubová *et al*., 2020). Since in this study we aimed to analyse how PAW treatment may impact plant-microbe symbiotic interactions in the rhizosphere - in particular on the AM symbiosis - the leguminous plant *L. japonicus* was used as plant experimental system, being able, unlike *A. thaliana*, to establish symbioses with both AM fungi and rhizobia.

To check if Ca^2+^-mediated perception of PAW is a plant general feature, we transformed *L. japonicus* with constructs encoding the bioluminescent Ca^2+^ reporter aequorin targeted to either the cytosol or the nucleus (Binci et al., 2024). Root segments from *L. japonicus* composite plants were challenged with freshly prepared PAW, tested at different dilutions. PAW was generated by exposing 50 ml deionized H_2_O for 5 min to a plasma torch operating at 900 W. The Ca^2+^ response of *L. japonicus* roots to treatment with PAW was monitored, by tracking changes in cytosolic ([Ca^2+^]_cyt_) and nuclear ([Ca^2+^]_nuc_) Ca^2+^ concentrations. As shown in Figure 1A-C, the treatment of *L. japonicus* roots with PAW (final dilution 1:2) triggered a fast and sustained elevation in [Ca^2+^]_cyt_, similarly to what previously observed in *A. thaliana* (Cortese *et al.,* 2021). No Ca^2+^ increases were detected in control samples, in which deionized H_2_O (without activation by plasma) was administered to *L. japonicus* roots. The magnitude of the induced Ca^2+^ signals, in terms of both [Ca^2+^] peak value (Fig. 1B) and integrated [Ca^2+^]dynamics (Fig. 1C), were found to correlate with the degree of PAW dilution, demonstrating a dose-dependent effect in the PAW-induced cytosolic Ca^2+^ changes triggered in *L. japonicus* roots. Taken together, these data confirm and extend our previous findings concerning Ca^2+^-mediated sensing of PAW by plants.

**Fig. 1.**
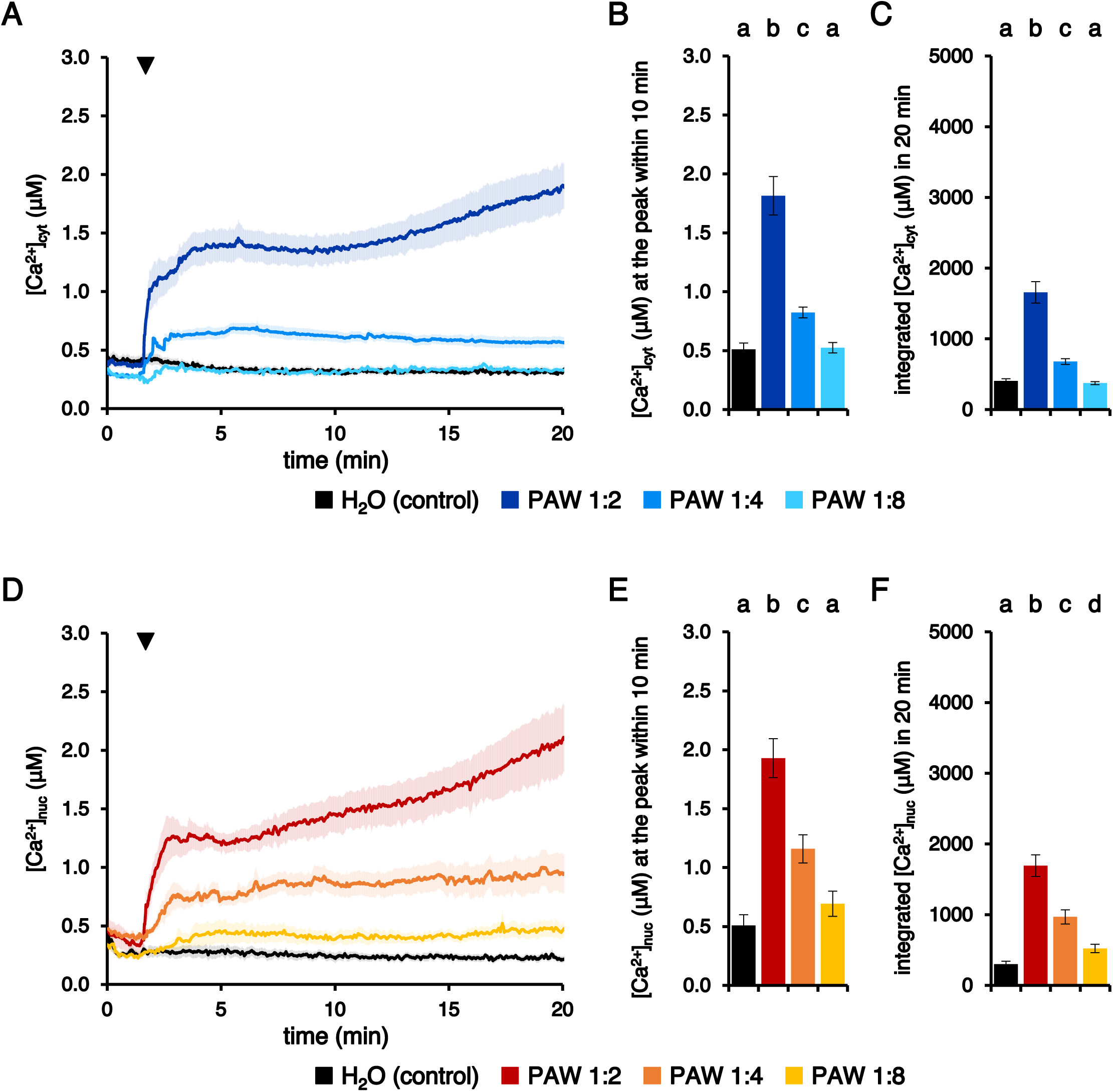
Monitoring dynamic changes in cytosolic (A-B) and nuclear (C-D) Ca^2+^ concentrations ([Ca^2+^]_cyt_ and [Ca^2+^]_nuc_) induced by plasma-activated water (PAW) in *L. japonicus* roots. *L. japonicus* roots expressing aequorin-based Ca^2+^ probes targeted to either the cytosol (A-B) or nucleus (C-D) were challenged with progressive dilutions of PAW (lighter colours indicate more diluted PAWs) generated by exposing deionized H_2_O to plasma torch-derived atmospheric plasma for 5 min at 900 W. Control samples were subjected to untreated deionized H2O (grey). Data are the means (solid lines) ± SE (shading) of ≥ 9 independent experiments. The arrowhead indicates the time of stimulus application (at 100 s). Statistical analyses were performed on the [Ca^2+^]_cyt_ peaks recorded in the first 10 min (B, E) and on the integrated [Ca^2+^]_cyt_ dynamics over 20 min (C, F). Bars labelled with different letters differ significantly (*p* < 0.05, Student’s *t* test).

To analyse if the treatment with PAW might induce changes in [Ca^2+^] not only in the cytosol, but also in the nucleus, PAW was administered to *L. japonicus* roots transformed with the construct encoding nuclear-targeted aequorin. As illustrated in Figure 1D-F, PAW was found to trigger significant elevations in [Ca^2+^]_nuc_, characterized by an amplitude correlated with PAW dilution. The Ca^2+^ response triggered by PAW 1:8, measured as integrated Ca^2+^ dynamics over 20 min, was significantly higher than the control (Fig. 1F), marking a difference from the pattern observed in the cytosol (Fig. 1C). This divergence adds a layer of complexity to our understanding of PAW-induced Ca^2+^ responses in plants and highlights for the first time the occurrence of nuclear Ca^2+^ changes triggered by PAW.

### 3.2 Plant cell viability is not affected by PAW treatment

Ca^2+^ measurement assays in *L. japonicus* demonstrated that PAW induced, at the highest concentration tested, sustained intracellular Ca^2+^ elevations, characterized by remarkable levels of Ca^2+^ (above 1 μM) in both cytosol and nucleus, at least during the first 20 min after administration of the stimulus (Fig. 1A, D). To check whether the long-lasting elevation in intracellular [Ca^2+^] induced by PAW may affect cell viability, the Evans blue test was applied on *L. japonicus* cell cultures after treatment with PAW. The viability of *L. japonicus* cells was found to remain unaltered following the application of PAW. Indeed, no significant increase in cell death was found either 1 h (Fig. 2A) or even 48 h (Fig. 2B) after PAW administration, in comparison with control samples. The absence of adverse effects on cell viability in response to PAW application validates its potential use as a benign intervention in plant-related applications.

**Fig. 2.**
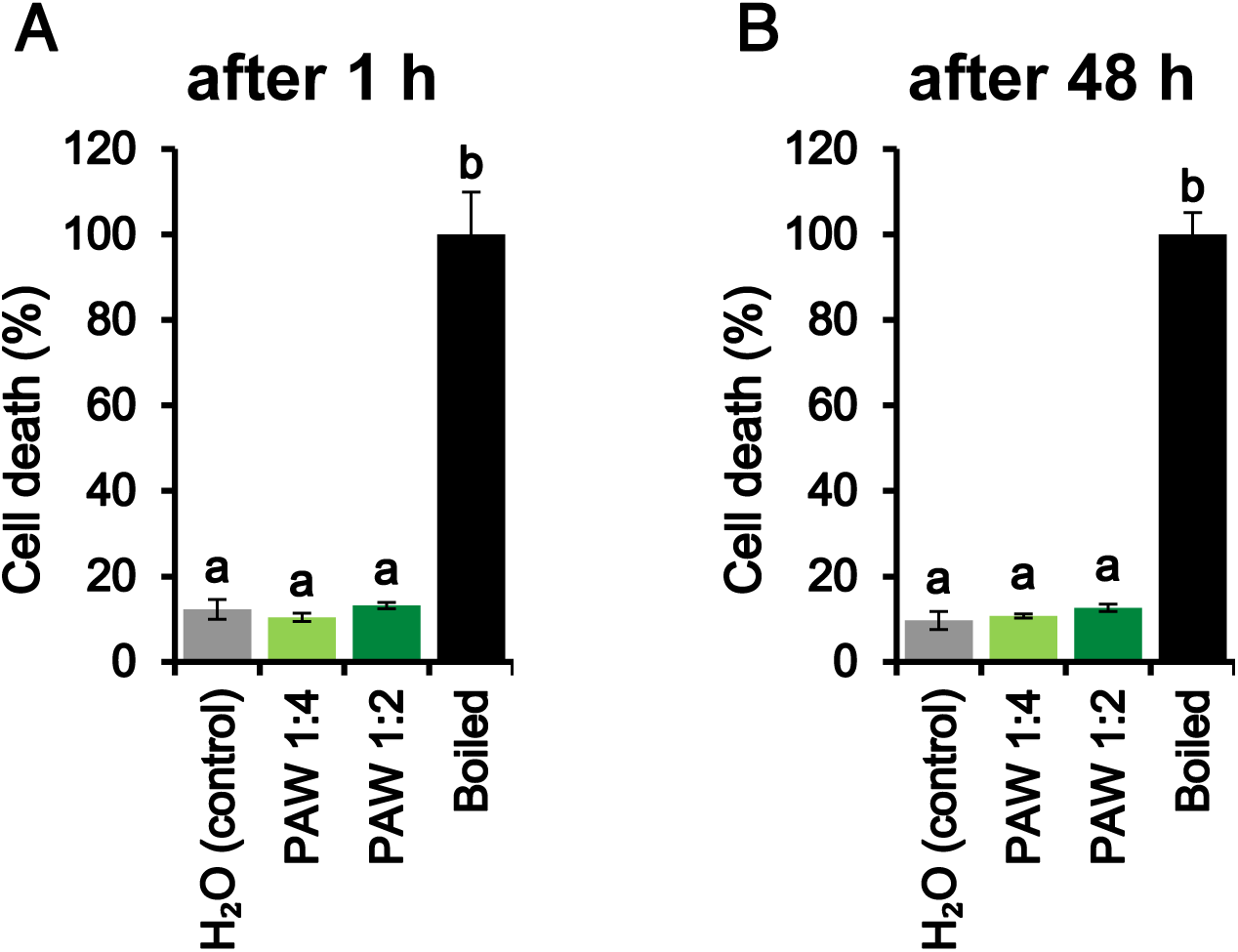
Viability of *L. japonicus* cell suspension cultures treated for either 1 h (A) or 48 h (B) with PAW generated by exposing deionized H_2_O to plasma torch-derived atmospheric plasma for 5 min at 900 W (1:4 diluted, light green; 1:2 diluted, dark green). Control cells were incubated with cell culture medium only (grey). The 100% value corresponds to cells treated for 10 min at 100°C (black bars). Bars labelled with different letters differ significantly (*p* < 0.05, Student’s *t* test).

### 3.3 Assessing the effects of PAW irrigation on the establishment of AM symbiosis between L. japonicus and the AM fungus R. irregularis

To evaluate the impact of the irrigation with PAW in terms of the establishment and development of the AM symbiosis of *L. japonicus* with *R. irregularis*, 14-day-old *L. japonicus* seedlings were co-cultivated with propagules of the AM fungus *R. irregularis* for 4 weeks. Plants were weekly irrigated alternating low-phosphate (20 μM Pi) Long Ashton liquid medium, to boost symbiosis establishment, to PAW 5’ treatments, according to the experimental set-up depicted in Supplementary Fig. S2. Different concentrations of PAW 5’ were tested (undiluted, 1:2, 1:4), whereas deionized water was used as a control. Figure 3 shows the quantification of the root colonization, according to the Trouvelot method (Trouvelot *et al.,* 1986) (Fig. 3A) and representative images obtained by light microscopy observations of the root apparatus of *L. japonicus* plants after 4 weeks of the various PAW treatments, in comparison with plants irrigated with H_2_O only (Fig. 3B). The obtained data revealed significant differences between PAW-treated plants and the controls. Treatment of *L. japonicus* plants with undiluted PAW 5’ resulted in a significant increase in all the symbiotic parameters examined, *i.e.* the frequency of mycorrhiza in the root system (F%), the intensity of mycorrhizal colonization (M%), and the abundance of arbuscules (A%). Moreover, the presence of vesicles was significantly higher in plants treated with PAW in comparison with control plants. On the other hand, no significant differences were observed when plants were irrigated with diluted (either 1:2 or 1:4) PAW (Fig. 3A). These data demonstrate that PAW positively modulates the establishment of AM symbiosis at an early phase of fungal colonization.

**Fig. 3.**
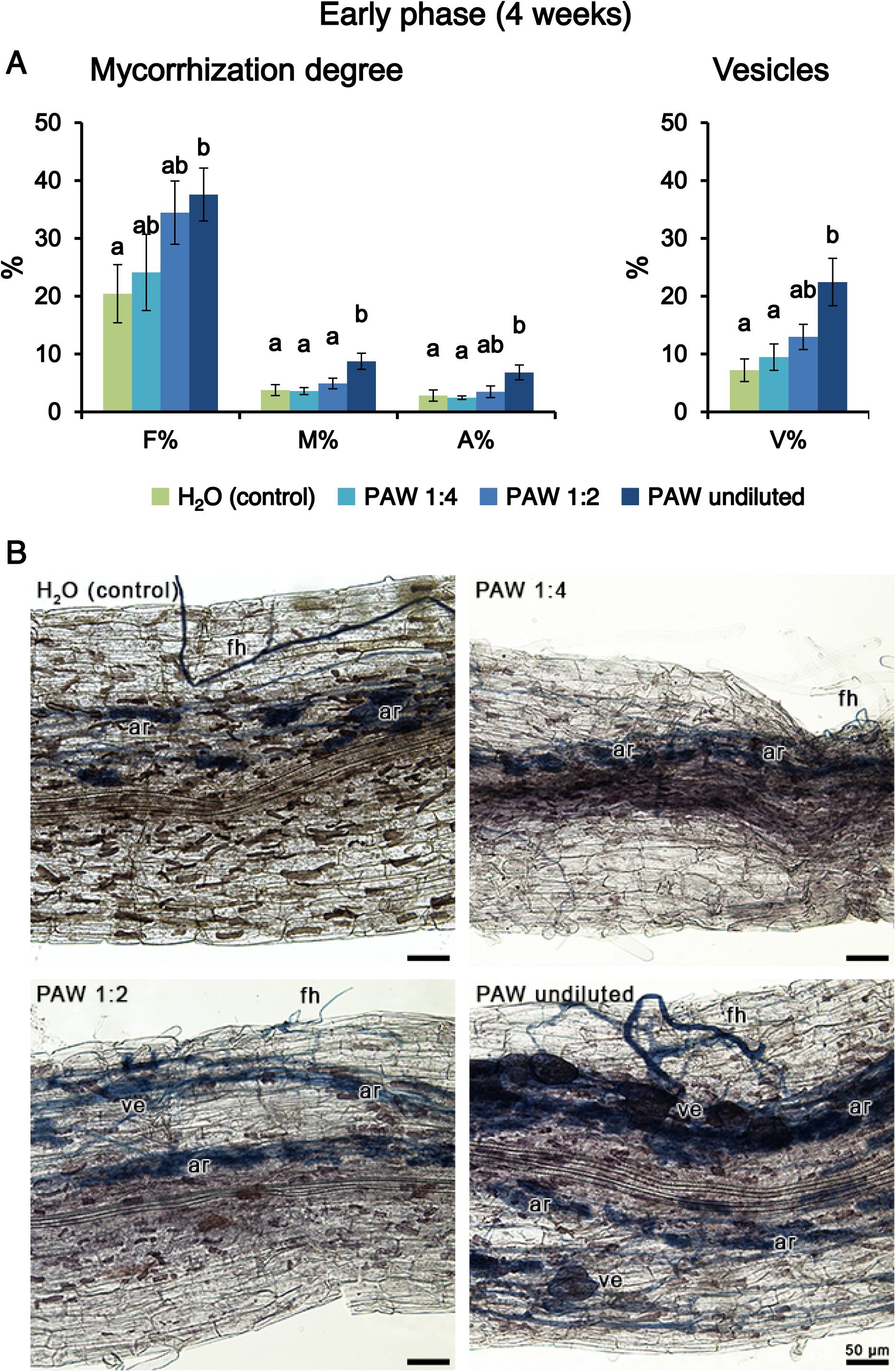
Phenotypic evaluation of mycorrhizal colonization after 4 weeks (early phase) in *L. japonicus* plants inoculated with the AM fungus *R. irregularis* and irrigated with PAW (1:4 diluted, light blue; 1:2 diluted, blue; undiluted, dark blue) or H_2_O only (control, green). PAW was obtained by exposing deionized H_2_O for 5 min to atmospheric plasma (PAW 5’) generated by a plasma torch operating at 900 W. A) The quantification of the degree of mycorrhization was conducted via the Trouvelot method. F%, frequency of mycorrhiza; M%, intensity of the mycorrhizal colonization; A%, arbuscule abundance. Vesicle abundance (V%) was also determined. Data are the means ± SE of 5 biological replicates from 2 independent experiments. Bars labelled with different letters differ significantly (*p* < 0.05, Student’s *t* test). B) Representative images of the colonized root apparatus of *L. japonicus* plants from the different condition groups. Fungal structures are highlighted in blue by ink-lactic acid staining. ar, arbuscule; fg, fungal hypha; ve, vesicle.

Given the positive impact of PAW 5’ on mycorrhizal colonization, another set of experiments was carried out with extended time durations (5 and 7 weeks post-inoculum) and including also a more intense PAW, *i.e.* H_2_O exposed to the plasma for 10 min (PAW 10’). Figure 4A shows that after 5 weeks there were no significant differences between PAW-treated plants and controls, with the only exception of the statistically significant variation in the frequency of mycorrhiza, where the PAW 10’ showed a negative effect on the root colonization level. It has to be noted that also the other symbiotic parameters in the PAW 10’ treatment showed a slight reduction trend, although not statistically significant. These results collectively contribute to a fine understanding of the impact of PAW treatments on the delicate balance of the AM symbiotic relationship between the AM fungus *R. irregularis* and *L. japonicus* plants during the mid-phase (5 weeks) of the experimental timeline. The analysis conducted at the late-phase time point (7 weeks) did not reveal significant differences among the various treatment groups, suggesting a recovery phase after the potentially detrimental effect exerted by PAW 10’ at the mid-time point. Indeed, all the plant root samples displayed an extremely high level of AM fungal colonization (close to 100% of the root segments), suggesting that saturation was reached (Fig. 4B). Therefore, the beneficial effects played by PAW 5’ seem to characterize the early phase of AM interaction, by promoting a faster fungal colonization of the plant root apparatus (Fig. 3), whereas when the symbiotic interaction between the partners is fully established there are no more visible differences in the symbiotic phenotype (Fig. 4).

**Fig. 4.**
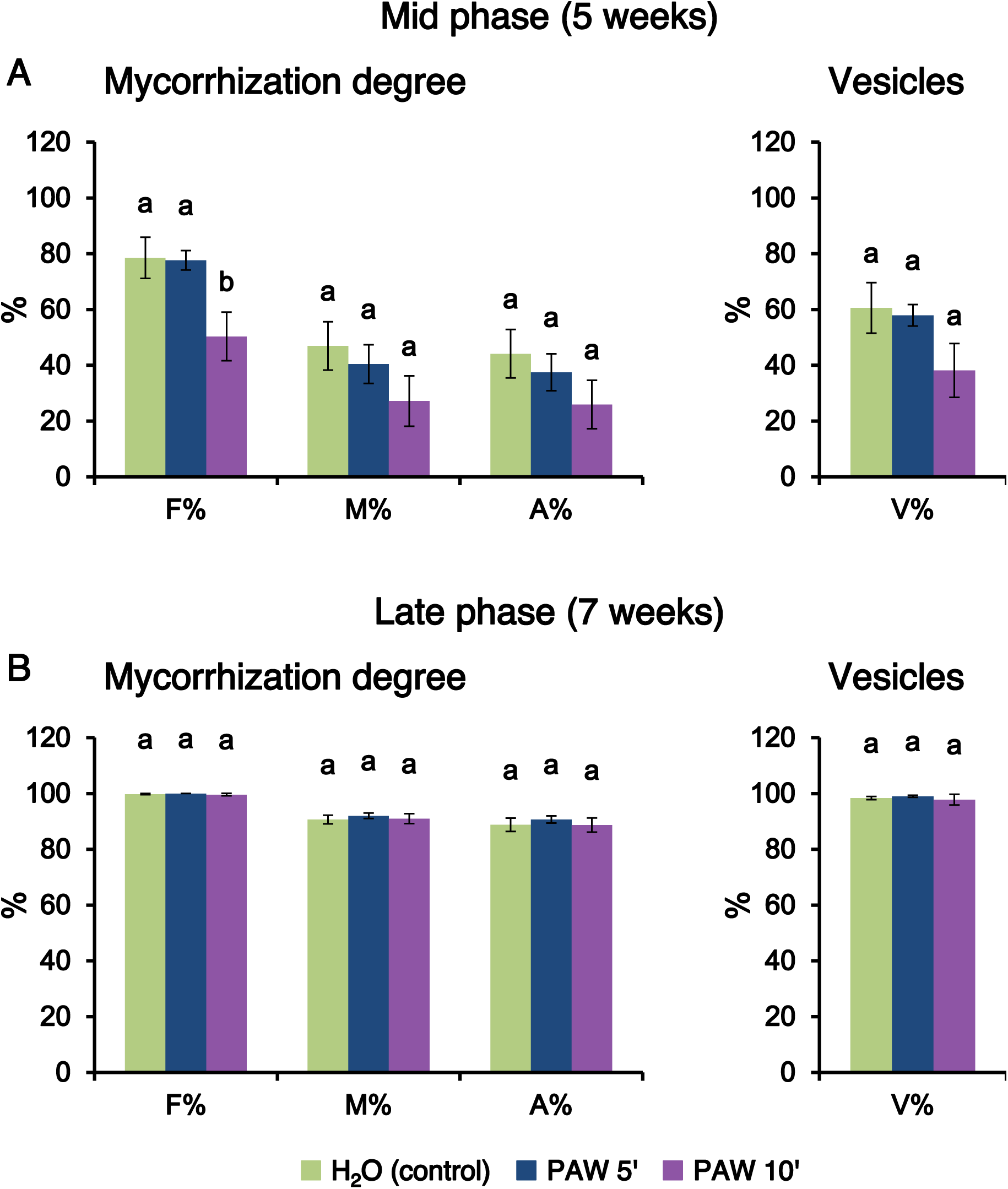
Phenotypic evaluation of mycorrhizal colonization after 5 weeks (A, mid phase) or 7 weeks (B, late phase) in *L. japonicus* plants irrigated with PAW 5’ (blue), PAW 10’ (violet) or H_2_O only (control, green). PAW was obtained by exposing deionized H_2_O for either 5 min (PAW 5’) or 10 min (PAW 10’) to atmospheric plasma generated by a plasma torch operating at 900 W. The quantification of the degree of mycorrhization was conducted via the Trouvelot method. F%, frequency of mycorrhiza; M%, intensity of the mycorrhizal colonization; A%, arbuscule abundance. Vesicle abundance (V%) was also determined. Data are the means ± SE of 5 biological replicates from 2 independent experiments. Bars labelled with different letters differ significantly (*p* < 0.05, Student’s *t* test).

### 3.4 Fresh weight analysis aligns with the root colonization levels

For each above-mentioned experimental conditions, measurements of fresh weight were carried out for both shoot and root at the different timepoints. At the early time point (4 weeks) a positive effect of PAW 5’ treatment was observed concerning both shoot and root biomasses in comparison with control (Fig. 5A). On the other hand, no statistically significant differences were observed in samples undergoing PAW 10’ treatment. At the mid time point (5 weeks), PAW 10’ exhibited a slight inhibitory impact on the fresh weight of the samples, indicative of a negative influence of PAW 10’ on the overall growth of the plants at this specific time interval. Samples treated with PAW 5’ did not show significant differences in either shoot or root fresh weight at mid phase (Fig. 5B). When the investigations were extended to the late time point of 7 weeks no significant differences in fresh weight were observed among the three treatments - control, PAW 5’, and PAW 10’-suggesting a certain degree of uniformity in growth responses at this later stage (Fig. 5C). These data confirm and extend the findings obtained by Trouvelot analyses (Fig. 3 and Fig. 4). To sum up, the obtained biomass data underscore the nuanced temporal dynamics of the effects of different PAWs on plant growth. Whereas a significant promoting effect is observed in PAW 5’-treated plants at the early phase (4 weeks), a mild depressive effect is detected with PAW 10’ treatment at the mid-time point (5 weeks). Both these divergent effects on plant biomasses appear to wane as the plants progress to the late time point of 7 weeks.

**Fig. 5.**
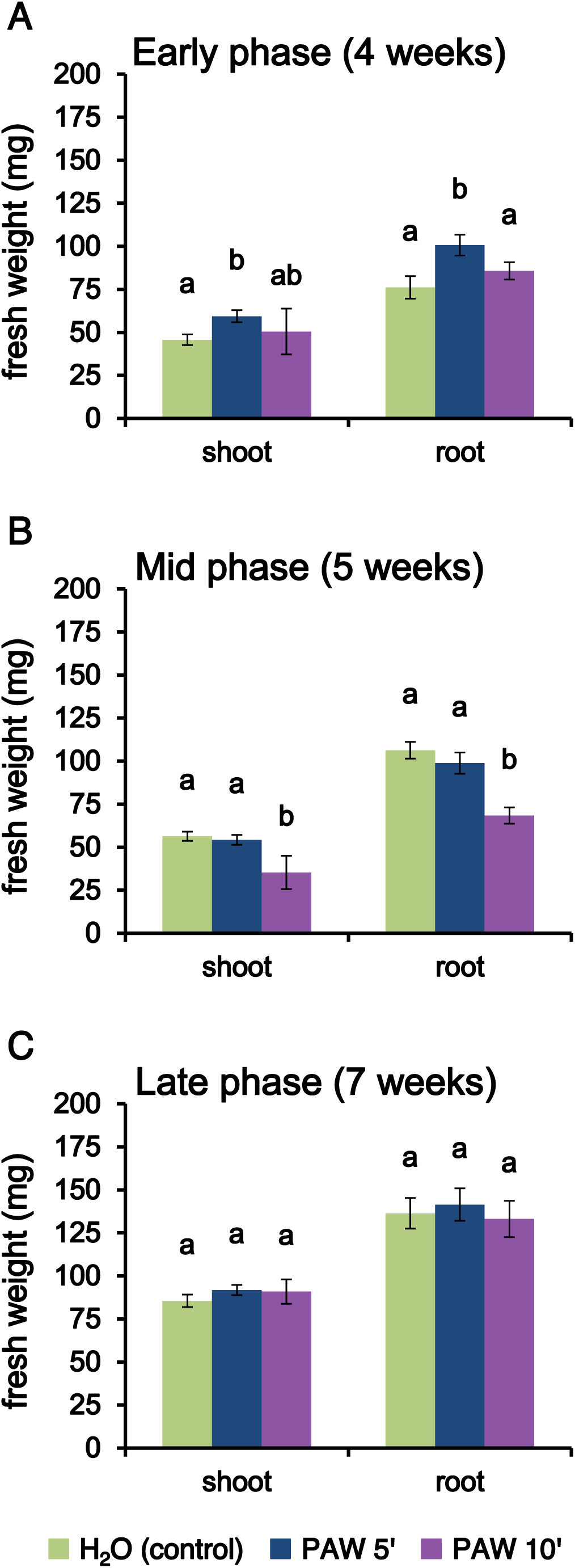
*L. japonicus* shoot and root biomass after repeated irrigations with PAW 5’, PAW 10’ or H_2_O only. Pots containing *L. japonicus* seedlings were treated weekly with either PAW 5’ (blue), PAW 10’ (violet) or H_2_O only (control, green) and harvested after 4 weeks (A, early phase), 5 weeks (B, mid phase) or 7 weeks (C, late phase) after the first treatment, respectively. PAW was obtained by exposing deionized H_2_O for either 5 min or 10 min to atmospheric plasma generated by a plasma torch operating at 900 W. Data are the means ± SE of 5 biological replicates from 2 independent experiments. Bars labelled with different letters differ significantly (*p* < 0.05, Student’s *t* test).

### 3.5 Inorganic phosphate analyses provide further insights into the modulatory effect played by PAW on AM symbiosis

The analysis of inorganic phosphate levels at different time points provides valuable insights into the effects of PAW treatment on the mineral nutrition of *L. japonicus* mediated by the AM fungus. Soluble inorganic phosphate was extracted from the leaf samples and quantified with a colorimetric assay based on the Malachite green dye. At the early phase and mid-phase time points (4 and 5 weeks), no statistically significant differences were observed among the treatments (Fig. 6). Even if small variations in the root colonization could be observed in treated plants with PAW 5’ (Fig. 3) and PAW 10’ (Fig. 4), they did not reflect on the accumulation of phosphate in the leaves of the host plant, possibly because of the limited time interval analysed (4-5 weeks). At the late phase time point (7 weeks), noteworthy variations emerged; specifically, samples treated with PAW 5’ exhibited a significantly higher concentration of leaf inorganic phosphate (Fig. 6). This observation points towards an improvement in mineral nutrition facilitated by PAW 5’, which might have stimulated phosphate uptake via the AM fungus or its assimilation processes.

**Fig. 6.**
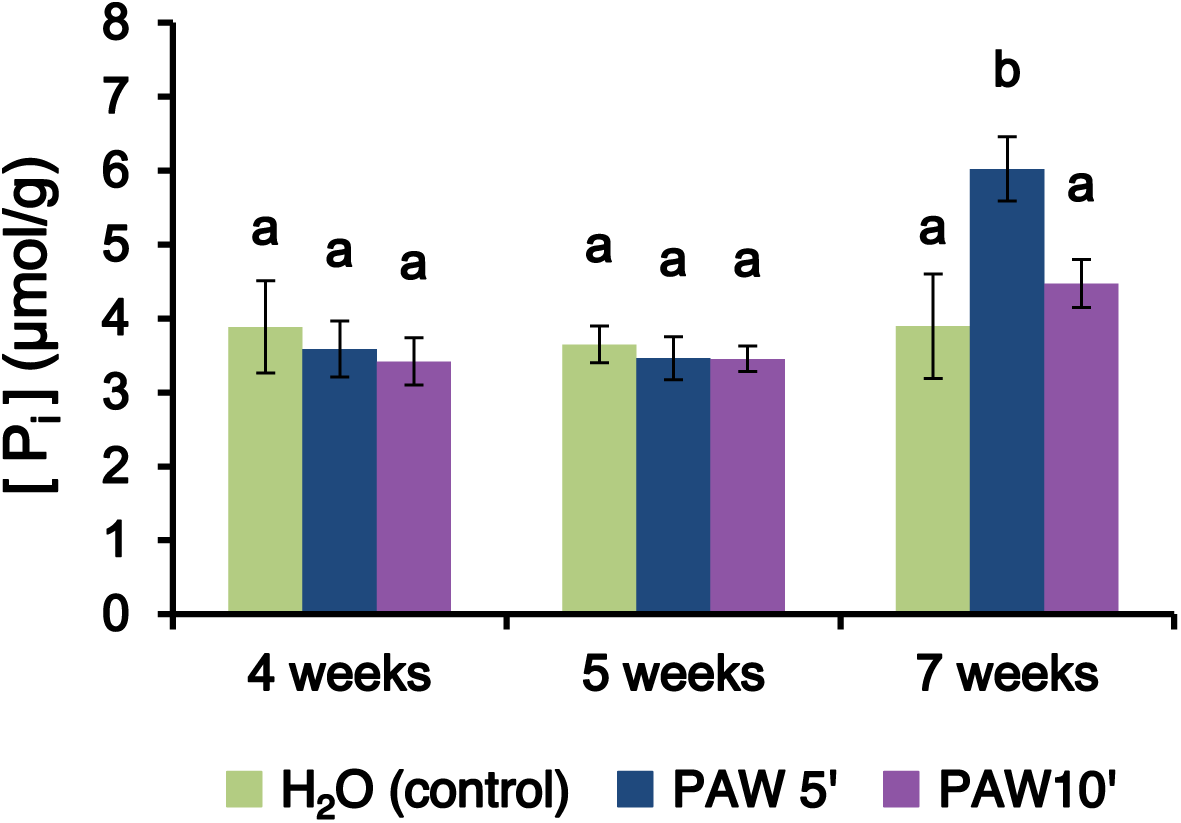
Inorganic phosphate (Pi) content in leaves of *L. japonicus* seedlings after repeated irrigations with PAW 5’, PAW 10’ or H_2_O only. Pots containing *L. japonicus* seedlings were treated weekly with either PAW 5’ (blue), PAW 10’ (violet) or H_2_O only (control, green) and harvested after 4 (A, early phase), 5 (B, mid phase) or 7 weeks (C, late phase) after the first treatment, respectively. PAW was obtained by exposing deionized H_2_O for either 5 min or 10 min to atmospheric plasma generated by a plasma torch operating at 900 W. Data are the means ± SE of 5 biological replicates from 2 independent experiments. Bars labelled with different letters differ significantly (*p* < 0.05, Student’s *t* test).

### 3.6 Nitrogen, carbon and sulphur analyses in L. japonicus shoot samples

Total nitrogen, carbon and sulphur concentrations within the aerial parts of the mycorrhizal *L. japonicus* plants collected at different time points after the onset of irrigations with either PAW 5’, PAW 10’, or H_2_O only were determined (Fig. 7). At 4 weeks PAW 5’ treatment resulted in a nitrogen content significantly higher than the control. In contrast, the PAW 10’ treatment did not show a similar increase in N content during this early stage. As the experiment progressed into the mid phase, a shift in the relationship between PAW treatments and N content became apparent. Indeed, at 5 weeks the PAW 10’ treatment, which is associated with lower fresh weight (Fig. 6), shows the highest N content. This fact could be interpreted as smaller plants having more concentrated N content. As the experiment reached its late phase (7 weeks), both PAW treatments exhibited an increase in respect with the control, even if only PAW 5’ treatment is statistically significant. As a general trend, PAW treatments resulted in increased N accumulation in the shoots (Fig. 7A). In terms of carbon content (Fig. 7B), the data exhibit an almost uniform behaviour across treatments and time intervals analysed, in line with the constant role of this element in herbaceous plants of this type and age. The data about sulphur content (Fig. 7C) indicate that the most stressful situations (plants undergoing PAW 10’ treatments) are associated with a statistically significant S accumulation. This phenomenon may be indicative of the activation of plant defence strategies, potentially involving the synthesis of cysteine-rich proteins and the production of antioxidant compounds (Adhikari et al., 2019). Such responses align with well-documented plant defence strategies and overall alerting pathways (Savvides *et al*., 2016).

**Fig. 7.**
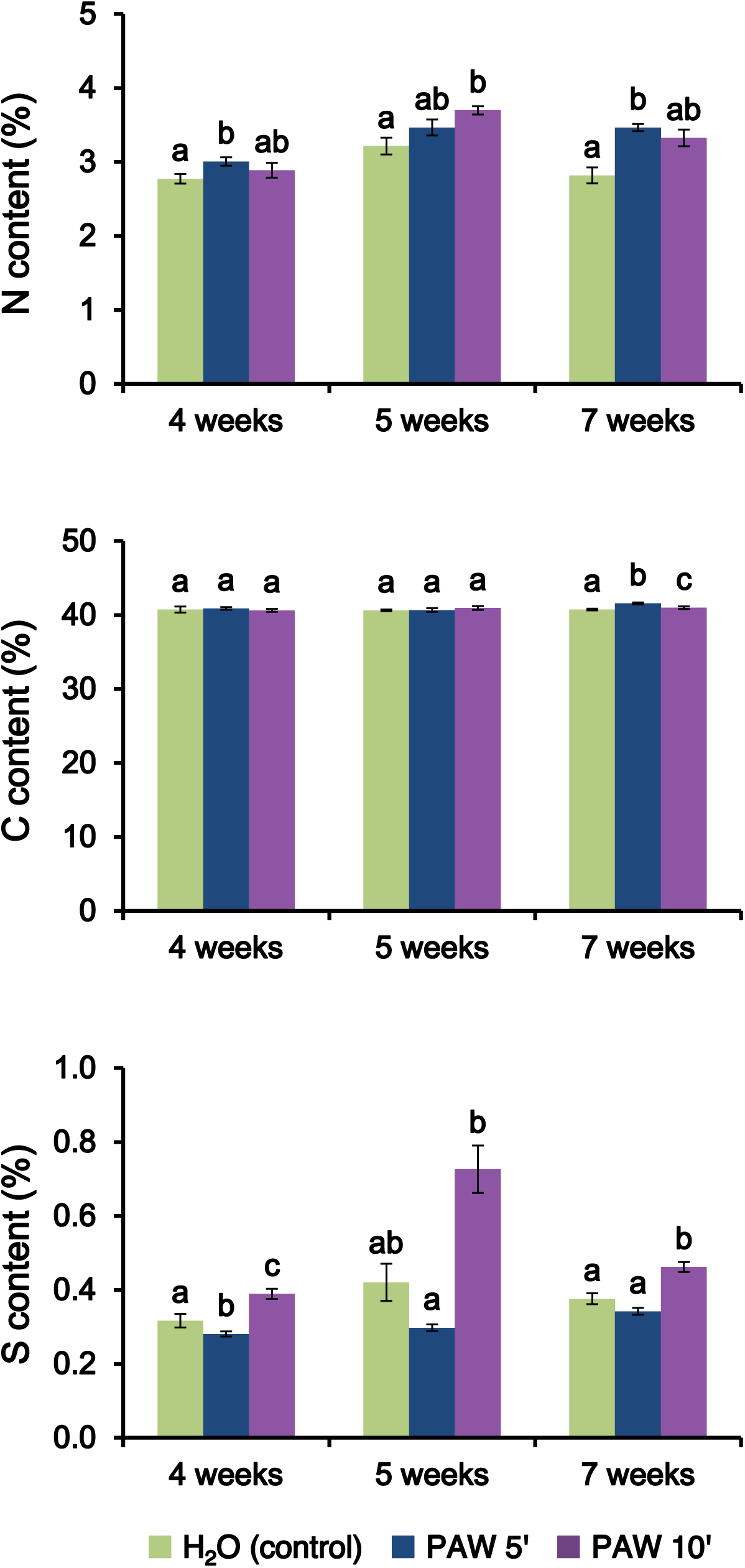
Quantification of nitrogen, carbon and sulphur content in dry shoots of *L. japonicus* seedlings after repeated irrigations with PAW 5’, PAW 10’ or H_2_O only. Pots containing *L. japonicus* seedlings were treated weekly with either PAW 5’ (blue), PAW 10’ (violet) or H_2_O only (control, green) and harvested after 4 (early phase), 5 (mid phase) or 7 weeks (late phase) after the first treatment, respectively. PAW was obtained by exposing deionized H_2_O for either 5 min or 10 min to atmospheric plasma generated by a plasma torch operating at 900 W. Data are the means ± SE of 5 biological replicates from 2 independent experiments. Bars labelled with different letters differ significantly (*p* < 0.05, Student’s *t* test).

## Discussion

Plasma-activated water (PAW) represents a novel and emerging eco-friendly technology with potential broad impact in agriculture, regarding plant fertilization, increased tolerance to abiotic stresses and control of pests (Holubová et *al.*, 2020; Guo et al., 2021; Gao et al., 2022). Despite in the last few years there has been a surge of papers in this field, information on how PAW and in general plasma treatments may impact plant-microbe symbiotic interactions is still scarce and only limited to nodulation (Pérez-Pizá et al., 2020; Abeysingha et al., 2024). In this study, we conducted an experimental investigation into PAW sensing mechanisms and effects on the initiation and progression of the mutualistic symbiosis between the model legume *L. japonicus* and the arbuscular mycorrhizal (AM) fungus *R. irregularis*.

Since PAW perception has recently been demonstrated to occur through Ca^2+^-mediated signalling in *Arabidopsis thaliana* (Cortese *et al*., 2021 and 2022), in this work we monitored early responses to different dilutions of PAW in *L. japonicus* roots expressing the bioluminescent Ca^2+^ reporter aequorin. Our results demonstrate that PAW triggers fast and sustained Ca^2+^ elevations not only in the cytosol, but also in the nucleus of *L. japonicus* root cells, adding a further layer of complexity in PAW-activated signal transduction pathways. Very recently, non-thermal plasma irradiation has been shown to induce a rapid increase in cytosolic Ca^2+^ in the liverwort *Marchantia polymorpha* expressing the GFP-based Ca^2+^ indicator GCaMP6f (Tsuboyama et al., 2024). These findings indicate that fast and transient intracellular Ca^2+^ elevations can be considered as hallmarks of initial cellular responses to both PAW and direct plasma treatments in plants.

Ca^2+^ measurements were used as an early assay to define the PAW doses to use in the following AM symbiosis experiments. Moreover, cell viability assays highlighted the non-toxic effect of PAW on plant cells, contributing to the growing body of evidence supporting the safety profile of PAW and its viability as an eco-friendly approach to enhance plant health and productivity (Pańka et al., 2022; Konchekov et al., 2023).

Our experimental framework involved the inoculation of *L. japonicus* seedlings with the AM fungus *R. irregularis*, followed by weekly irrigation with PAWs, generated by exposing H_2_O to the plasma for either 5 min (PAW 5’) or 10 min (PAW 10’), for different time intervals, and the following analyses of mycorrhizal fungal colonization after 4, 5 and 7 weeks. The obtained results indicate that the treatments with PAW shape AM development and functioning, in a manner dependent on the time interval of water exposure to the plasma and on the duration of plant irrigation with PAW. Our investigation reveals that, while the administration of PAW 5’ displays a significantly promoting effect on AM symbiosis, by accelerating the establishment of the *L. japonicus - R. irregularis* symbiotic relationship, PAW 10’ seems to exert a mild depressive effect, at least in the earliest phase of the plant-microbe interaction. An intriguing possibility is that a longer exposure of water to the plasma, and the consequent higher content of ROS and RNS in the PAW, may activate a pre-alert state in the plant, mounting a response more directed to the activation of defence mechanisms against environmental stresses, such as potential pathogens (Adhikari *et al*., 2020), rather than the accommodation of the fungal symbiont.

The determination of inorganic phosphate levels in the leaves and the chemical analysis of N, C, S content in the entire shoots of PAW-treated plants *versus* control plants allowed to link PAW composition and AM-mediated plant nutrient uptake. PAWs generated by either plasma torches or dielectric barrier discharge sources are enriched in H_2_O_2_, nitrate (NO_3_ ^-^) and nitrite (NO_2_ ^-^) (Cortese et al., 2021 and 2022). H_2_O_2_ is a well-known ROS that plays a dual role in plant biology, acting as both a stress inducer and a signalling molecule (Smirnoff & Arnaud, 2019). Nitric oxide (NO), a signalling molecule derived from NO_3_^-^ and NO ^-^ has multifaceted roles in plants, including mediation of root growth and modulation of stress responses (Besson-Bard *et al.,* 2008a). Moreover, NO has been demonstrated to play a regulatory role during plant interactions with both pathogenic and mutualistic fungi (Calcagno et al., 2012; Martínez-Medina *et al*., 2019). In both NO- and ROS-based signal transduction pathways, Ca^2+^ has been proposed to act as intracellular mediator (Besson-Bard *et al*., 2008b; Gilroy *et al*., 2014 and 2016), with mutual interplay resulting in the amplification of the respective signalling mechanisms (Marcec *et al*., 2019). Considering the relatively high concentration of NO_3_^-^ and NO_2_^-^ in plasma torch-generated PAW (Cortese et al., 2021), it is likely that treatment with PAW provides the plant with a nutritional advantage, along with the putative activation of H_2_O_2_- and NO-triggered signalling pathways, leading to either a priming status or facilitating the accommodation of the symbiotic fungus. Further investigations are needed to address the importance of these two effects of treatment with PAW on plant physiology and interactions with beneficial microbes in the rhizosphere.

The data obtained in this work concerning the effects induced by the application of PAW on AM symbiosis in *L. japonicus* align well with the model proposed by Song *et al*. (2020) about the effects of plasma treatments on plant responses, that critically depend on the operational parameters of the plasma source. Indeed, the impact of PAW on AM symbiosis in *L. japonicus* can be explained with the intriguing concept of plant hormesis, which is a distinctive dose-response phenomenon characterized by stimulatory responses at low doses and inhibitory responses at higher doses (Agathokleous *et al*., 2019; Schirrmacher, 2021; Erofeeva, 2022 and 2024). Based on the obtained results, we propose a model about the hermetic effect of PAW treatment on AM symbiosis, which encompasses the following: i) PAW 5’: promotion of AM fungal colonization and plant growth in the early phase, along with an increase of phosphate uptake in the late phase; ii) PAW 10’: delay in AM fungal colonization and plant growth during early and mid-phases, but no long-lasting side effects in the late phase by (Fig. 8).

**Fig. 8.**
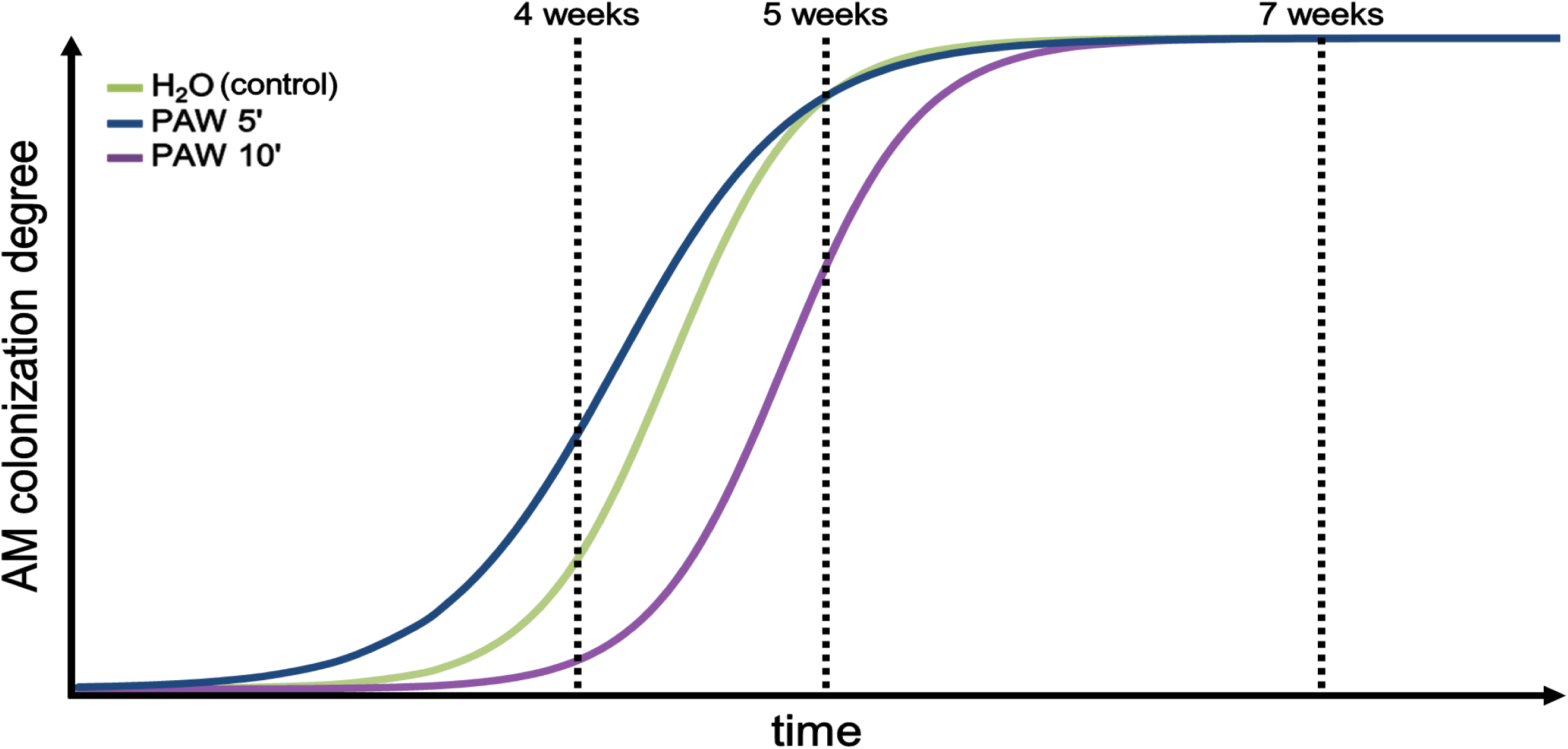
Proposed model of hormetic effect played by PAW on the establishment and development of the AM symbiosis.

## Conclusions

This study underscores the potential of PAW technology to finely tune the ability of plants to engage in symbiotic relationships with beneficial microorganisms of the rhizosphere, in particular AM fungi. As AM symbiosis is crucial to overall plant health and productivity, understanding how PAW may affect it is fundamental for large scale agricultural implementation of PAW. Our results suggest that plasma treatments may be used to improve plant mineral nutrition and/or induce a pre-alert state (priming) through a controlled generation and administration of PAW to plants. Further investigations are needed to analyse the fine balance, under different PAW treatment regimes, between plant accommodation responses towards beneficial microbes and plant defence responses against potential microbes.

## CRediT authorship contribution statement

**Enrico Cortese:** Investigation, Formal analysis, Visualization, Writing-original draft. **Filippo Binci:** Investigation, Formal analysis, Writing-original draft. **Erfan Nouri:** Investigation. **Arianna Capparotto:** Investigation, Formal analysis. **Alessio G. Settimi:** Methodology. **Manuele Dabalà**: Resources, Methodology. **Vanni Antoni:** Methodology, Writing - review and editing. **Andrea Squartini:** Methodology, Writing - review and editing, Funding acquisition. **Marco Giovannetti:** Supervision, Methodology, Resources, Writing - review and editing, Funding acquisition. **Lorella Navazio:** Conceptualization, Supervision, Methodology, Resources, Project administration, Writing - original draft, Writing - review and editing, Funding acquisition.

## Acknowledgments

We thank the Plant Genome Editing facility of the Department of Biology, University of Padova (Italy) and the central Chemical Laboratory (La-Chi) of the Department of Agronomy, Food, Natural resources, Animals and Environment, University of Padova (Italy) for skilful technical support. This work was supported by grants from the Italian Ministry of University (PRIN 2022 - grant number 2022NW97JX to L.N.), from the University of Padova (Progetti di Ricerca Dipartimentali-PRID) (grant number BIRD214519 to M.G.) and from the European Union-NextGenerationEU (2021 STARS Grants@Unipd programme P-NICHE to M.G.).

## Supplementary materials

**Supplementary Fig. S1.** Representative image of the generation of plasma-activated water (PAW) by exposing water to a plasma torch.

**Supplementary Fig. S2.** Experimental set-up for the evaluation of the effect of PAW treatment on the co-cultivation of *L. japonicus* seedlings with the AM fungus *R. irregularis*.

