## Supplementary figures S1-S2 for "Modulatory effect of plasma-activated water on arbuscular mycorrhizal symbiosis in *Lotus japonicus*"

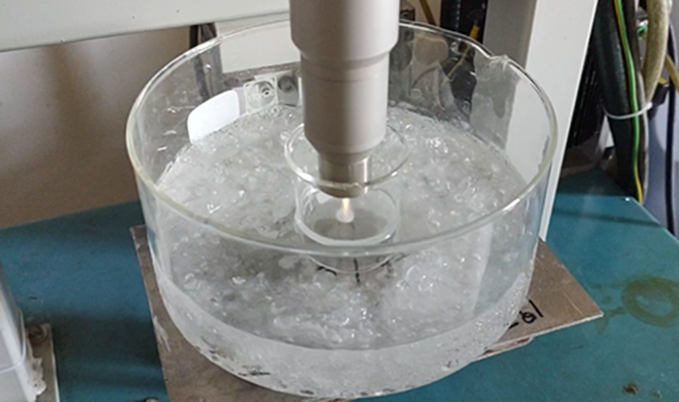

**Supplementary Figure S1.** Representative image of the generation of plasma-activated water (PAW) by exposing water to a plasma torch.

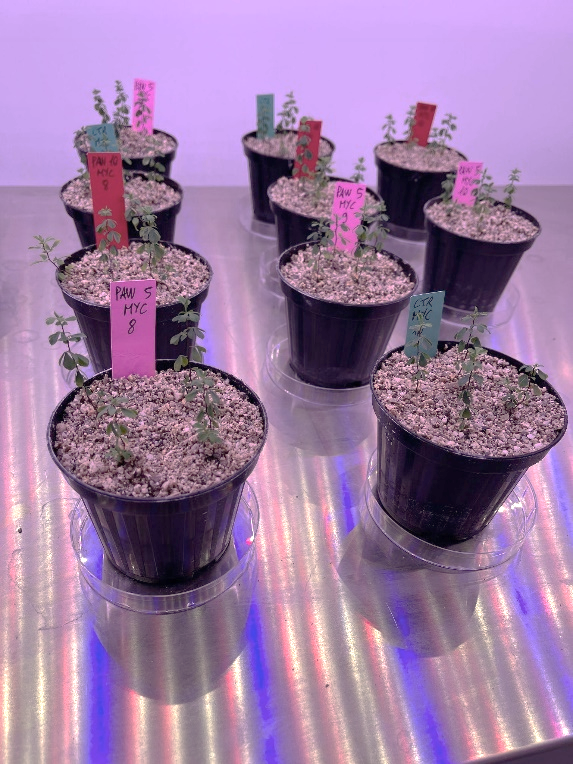

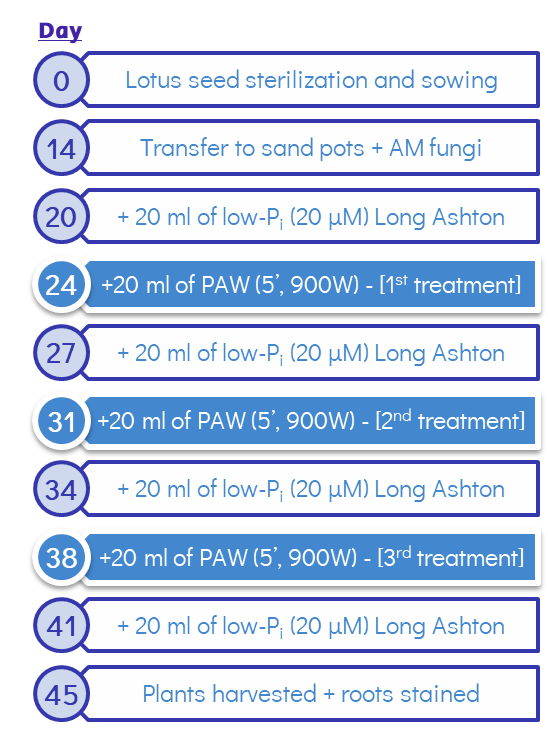

B

A

**Supplementary Figure S2.** Experimental set-up for the evaluation of the effect of PAW treatment on the co-cultivation of *L. japonicus* seedlings with the AM fungus *R. irregularis.* A) Scheme of the irrigation plan. B) Representative image of the randomized pots containing *L. japonicus* seedlings subjected to different treatments in the plant growth chamber.
